# Direct Imaging of Liquid Domains in Membranes by Cryo Electron Tomography

**DOI:** 10.1101/2020.02.05.935684

**Authors:** Caitlin E. Cornell, Alexander Mileant, Niket Thakkar, Kelly K. Lee, Sarah L. Keller

## Abstract

Images of micron-scale domains in lipid bilayers have provided the gold standard of model-free evidence to understand the domains’ shapes, sizes, and distributions. Corresponding techniques to directly and quantitatively assess smaller (nanoscale and submicron) liquid domains have been lacking, leading to an inability to answer key questions. For example, researchers commonly seek to correlate activities of membrane proteins with attributes of the domains in which they reside; doing so hinges on identification and characterization of membrane domains. Although some features of membrane domains can be probed by indirect methods, these methods are often constrained by the limitation that data must be analyzed in the context of models that require multiple assumptions or parameters. Here, we address this challenge by developing and testing two new methods of identifying submicron domains in biomimetic membranes. Both methods leverage cryo-electron tomograms of ternary membranes under native solution conditions. The first method is optimized for probe-free applications: domains are directly distinguished from the surrounding membrane by their thickness. This technique measures area fractions of domains with quantitative accuracy, in excellent agreement with known phase diagrams. The second method is optimized for applications in which a single label is deployed for imaging membranes by both high-resolution cryo-electron tomography and diffraction-limited optical microscopy. For this method, we test a panel of probes, find that a trimeric mCherry label performs best, and specify criteria for developing future high-performance, dual-use probes. These developments have led to the first direct and quantitative imaging of submicron membrane domains under native conditions.

**SIGNIFICANCE STATEMENT:** Fluorescence micrographs that capture the sizes, shapes, and distributions of liquid domains in model membranes have provided high standards of evidence to prove (and disprove) theories of how micron-scale domains form and grow. Corresponding theories about smaller domains have remained untested, partly because experimental methods of identifying submicron domains in vesicles under native solvent conditions have not been available. Here we introduce two such methods. Both leverage cryo-electron tomography to observe membrane features far smaller than the diffraction limit of light. The first method is probe-free and identifies differences in thicknesses between liquid domains and their surrounding membranes. The second method identifies membrane regions labeled by an electron-dense, fluorescent protein, which enables direct comparison of fluorescence micrographs with cryo-electron tomograms.

## INTRODUCTION

Seeing is believing, which makes images powerful. Current advances in microscopy have revolutionized our understanding of cellular components, macromolecular assemblies, protein structure, and membrane organization. For example, images of micron-scale synapses in stimulated immune cells have successfully led to the development of quantitative models of membrane protein interactions (1). Similarly, direct imaging has demonstrated that vacuole membranes in living yeast cells phase separate (2, 3) and that model and cell-derived membranes exhibit critical phenomena (4, 5). However, in all of these examples, the membrane features span micrometer length scales. Challenges persist in observing membrane features that are far smaller than the diffraction limit of light, especially in model lipid vesicles under native solution conditions. As a result, a wide range of quantitative questions has remained impossible to answer. For example, if a vesicle membrane contains nanodomains, what are the sizes and distributions of those domains across the vesicle surfaces? Similarly, do submicron domains fit quantitative predictions of modulated phases or of microemulsions (6, 7)?

New approaches are needed in order to overcome current limitations. Transmission electron microscopy (TEM) can achieve near-atomic resolution, and freeze-fracture TEM has successfully been used to identify coexisting solid and liquid phases in simple, lipid membranes (8–11). However, freeze-fracture is an unwieldy technique that images a metal-shadowed surface of a membrane. To date, freeze-fracture has achieved contrast between liquid domains and the rest of the membrane only when membranes contain large protein complexes (2). A more common way of identifying submicron liquid domains by TEM is gold-labeling of proteins and lipids (e.g. (12, 13)). This method results in over-counting (which can be misinterpreted as self-clustering proteins) if labeling uses both primary and secondary antibodies or if multiple labels are conjugated to a single antibody (14). Other methods of imaging submicron liquid domains have their own limitations. Atomic force microscopy (AFM) requires deposition of membranes on solid substrates, which typically captures domains in non-equilibrium sizes and shapes (15–21). Near-field scanning optical microscopy (NSOM) places cantilevers in contact with membranes, which may alter membrane structures (22). Standard super-resolution optical techniques cannot image small enough features, and expansion microscopy relies on cross-linked proteins (23).

Here, we introduce and test two new methods for identifying submicron domains in membranes from direct cryo-electron tomography (cryo-ET) images. To our knowledge, we (and the jointly-submitted manuscript by Heberle et al.) are the first to collect cryo-ET images of ternary model membranes under native solution conditions. Of the two new methods, the first is entirely label-free and leverages differences in thicknesses of the domains versus the rest of the membrane (Fig. 1). We benchmark this label-free approach against known phase diagrams to demonstrate that it accurately quantifies the area fractions of coexisting liquid-ordered (*L*_o_) and liquid-disordered (*L*_d_) phases in membranes. The second method employs a probe that is fluorescent, electron-dense, and labels the membrane through a single binding site. Our goal is for the probe to enable direct, model-free comparisons for a single vesicle sample analyzed by both fluorescence microscopy and electron microscopy. We test a panel of probes and find that a trimeric mCherry label performs best in this role.

**Figure 1.**
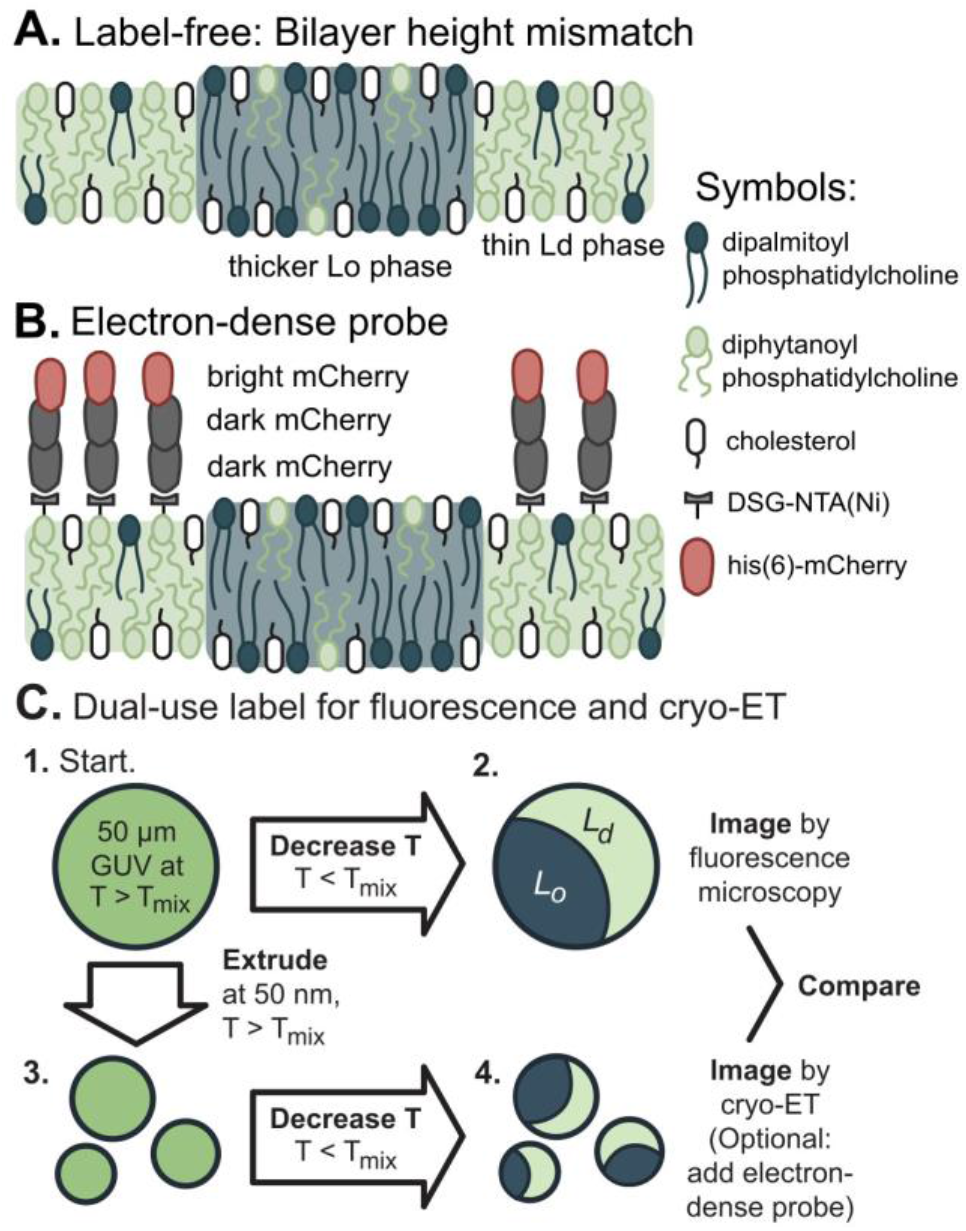
Two methods for direct identification of lipid domains in vesicles using cryo-ET. A. The label-free method identifies the difference in thickness between the *L*_o_ phase and *L*_d_ phase. B. The labeling method deploys a linear trimer of mCherry that is both fluorescent and electron-dense. A his6-tag on the terminal mCherry protein binds to a nickel-chelated lipid that preferentially partitions to the Ld phase. C. The mCherry label enables imaging a single starting solution of giant unilamellar vesicles (GUVs) by both fluorescence microscopy and cryo-ET.

## RESULTS

Figure 1 summarizes the two methods we developed for identifying coexisting liquid disordered (*L*_d_) and liquid ordered *(*L*_o_)* phases in vesicles by cryo-ET (Fig. 1). The first method exploits the difference in bilayer thicknesses of the phases. The second method employs a label that preferentially partitions to the *L*_d_ phase. We tested these methods on systems that represent the broad class of membranes known to separate into macroscopic *L*_d_ and *L*_o_ phases in GUVs. Specifically, we imaged uncharged membranes composed of ternary mixtures, consisting of a lipid with a low melting temperature (diphytanoyl-phosphocholine, DiPhyPC)), a lipid with a high melting temperature (dipalmitoyl-phosphocholine, DPPC), and a sterol (cholesterol) (24). Cryo-ET resolves submicron features of intact vesicles in aqueous environments. Therefore, to our knowledge, we (and the jointly-submitted manuscript by Heberle et al.) are the first to image submicron domains in membranes with coexisting liquid phases under native solvent conditions.

To maximize differences between the *L*_d_ and *L*_o_ phases so they are distinguishable, we mixed the lipids in ratios that fall along an unusually long tie-line (Fig. 2A and Table 1) (24). The endpoints of tie-lines represent the lipid compositions of the two phases. As a result, when we use cryo-ET to image submicron vesicles, we expect to observe *L*_o_ domains that are significantly (~1 nm) thicker than membranes of the surrounding *L*_d_ phase (20). The same concept applies when we use fluorescence microscopy to image GUVs labeled with Texas Red DHPE; *L*_o_ domains are significantly darker than the surrounding *L*_d_ phase (Fig. 2B and (24)).

**Figure 2.**
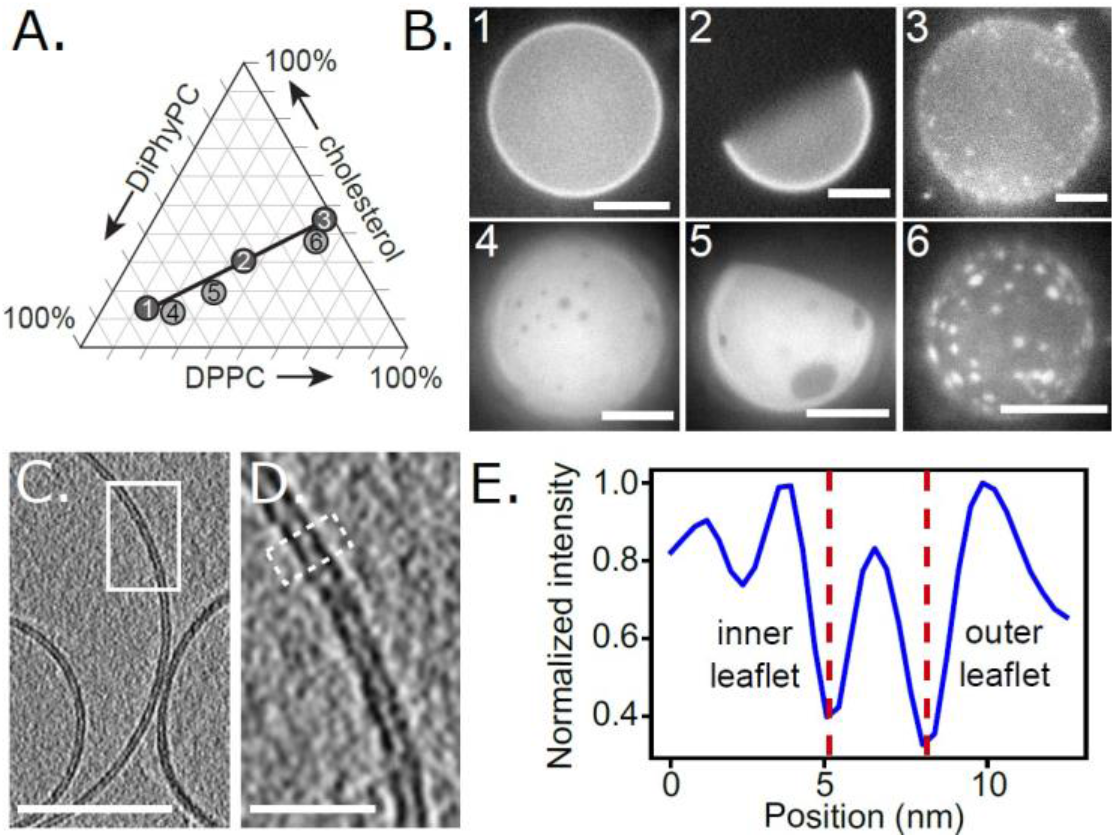
The fraction of membrane area in the *L*_o_ phase increases monotonically along tie-lines. **A**. All possible ternary mixtures of DiPhyPC, DPPC, and cholesterol fall within the triangle. Ratios 1, 2, and 3 lie on a tie-line at 22°C (24). Ratios 4, 5, and 6 lie on an extrapolated line parallel to the known tie-line. **B.** Representative fluorescence micrographs of GUVs made from Ratios 1-6. The GUVs contain 0.8 mol% of the dye Texas Red DHPE, which preferentially partitions to the *L*_d_ phase. To facilitate visualization of area fractions, images were captured shortly after domains nucleated, before all domains completely coalesced. **C**. Slice at 0° through a cryoET tomogram of a field of vesicles made from Ratio 4. Bilayer regions are resolvable as two distinct monolayer leaflets. Scale bar is 50 nm. **D**. Enlarged image of the area in the white box in Panel C. Scale bar is 10 nm. **E**. A linescan reveals two troughs, which correspond to the clearly resolved dark bands of the inner and outer leaflets of the membrane in Panel D. The linescan was 10 pixels wide and taken across the area in Panel D outlined in the white dashed line.

**Table 1:** Vesicles were produced from six lipid ratios. Ratios 1, 2, and 3 fall on a tie-line measured at 22°C from (24). Ratios 4, 5, and 6 fall on a parallel line. When trimeric mCherry probes were used, DGS-NTA(Ni) lipids were included in each mixture, replacing, 2 mol% of DiPhyPC lipids in Ratios 2 and 5. Values of the mole % of all lipids that are expected to be in the *L*_o_ phase are derived from two methods: fluorescence microscopy of GUVs and nuclear magnetic resonance (NMR) of multilamellar vesicles (24). Uncertainties in the mole % of *L*_o_ phase are ± 6%, propagated from NMR measurements of tie-line endpoints (24).

| | mole %<br>DiPhyPC | mole %<br>DPPC | mole %<br>Chol | mole %<br>$L_o$ phase |
| --- | --- | --- | --- | --- |
| Ratio 1 | 68 | 17 | 15 | 0% |
| Ratio 2 | 35 | 35 | 30 | 50% |
| Ratio 3 | 6 | 52 | 42 | 100% |
| Ratio 4 | 66 | 21 | 13 | 10% |
| Ratio 5 | 48 | 32 | 20 | 20% |
| Ratio 6 | 5 | 58 | 37 | 90% |

One of our central goals was to quantitatively benchmark cryo-ET results for 10-100 nm vesicles against fluorescence microscopy results for vesicles roughly a thousand times larger. To ensure that the lipid composition of vesicles did not vary with their size, we made careful choices about how we produced vesicles. Because different techniques incorporate different ratios of lipids into vesicles (25–27), we produced all vesicles by the same technique: electroformation (Fig. 1C). We maintained some of these vesicles as giant unilamellar vesicles (GUVs) to image them by fluorescence microscopy, and we extruded others through 50- or 100-nm pores to image by cryo-ET (Fig. 2 and SI Appendix, Fig. S1-S2).

To identify *L*_d_ and *L*_o_ domains by the first method, we took a probabilistic approach, separating distributions of bilayer thicknesses into component parts associated with each phase. Critically, this approach makes no assumptions about the size, spatial arrangement, or absolute thicknesses of domains. First, we collected cryo-electron tomograms for two types of control vesicles, which we made from lipid ratios that lay at the two ends of a tie-line (Ratios 1 and 3). To identify the two leaflets of vesicle membranes, we performed a Canny edge filter to the central slice of each tomogram. We established an objective procedure to determine apparent bilayer thicknesses; specifically, we evaluated the minimum distance from every pixel on the inner leaflet of vesicles to all possible pixels on the outer leaflet (Fig. 3A and Fig. 4). Because this procedure identifies differences in thicknesses rather than absolute thicknesses, it is robust to subtle changes in how different research groups might defocus tomograms, apply contrast transfer functions, or create edge filter algorithms.

**Figure 3.**
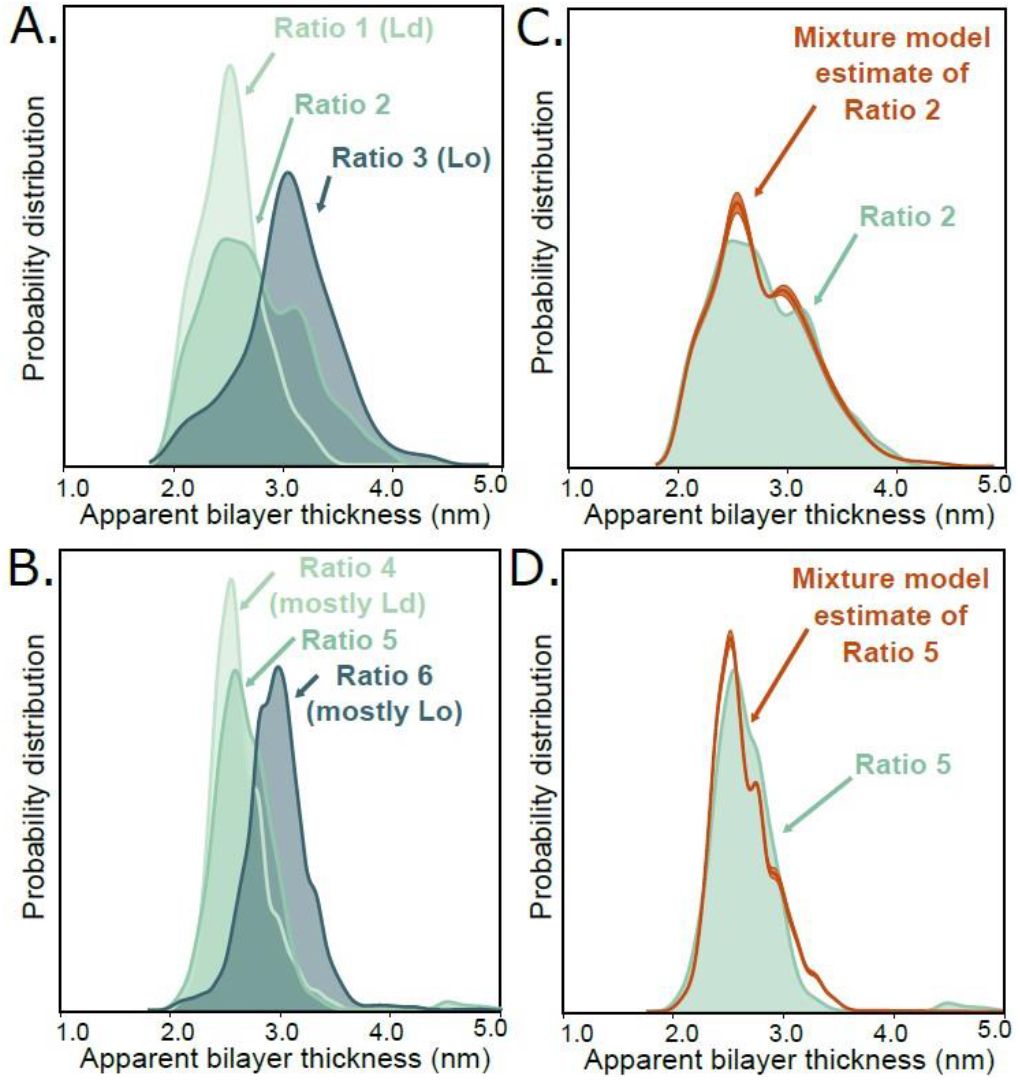
Quantitative agreement between ratios of *L*_o_ and *L*_d_ phases measured by cryo-ET and ratios expected from GUV phase diagrams. **A, B**. Approximately 5000 apparent bilayer thickness values are plotted for each lipid ratio (derived from measurements of the central slice of 20-30 vesicles, representing 10 tomograms of vesicle fields per lipid ratio). For all six ratios, Gaussian kernel density estimates (which plot the probability of measuring each distance, similar to a histogram) were calculated for all three ratios. **C, D**. The ratio of the membrane area in the *L*_o_ phase vs. the *L*_d_ phase can be estimated directly from images using a mixture of kernel density estimates, calculated using Ratios 1 and 3 for panel C and Ratios 4 and 6 for panel D. For Ratio 2 and Ratio 4, this procedure yields area ratios of 43:57 ± 3 *L*_d_:*L*_o_ and 83:17 ± 2 *L*_d_:*L*_o_, respectively.

**Figure 4.**
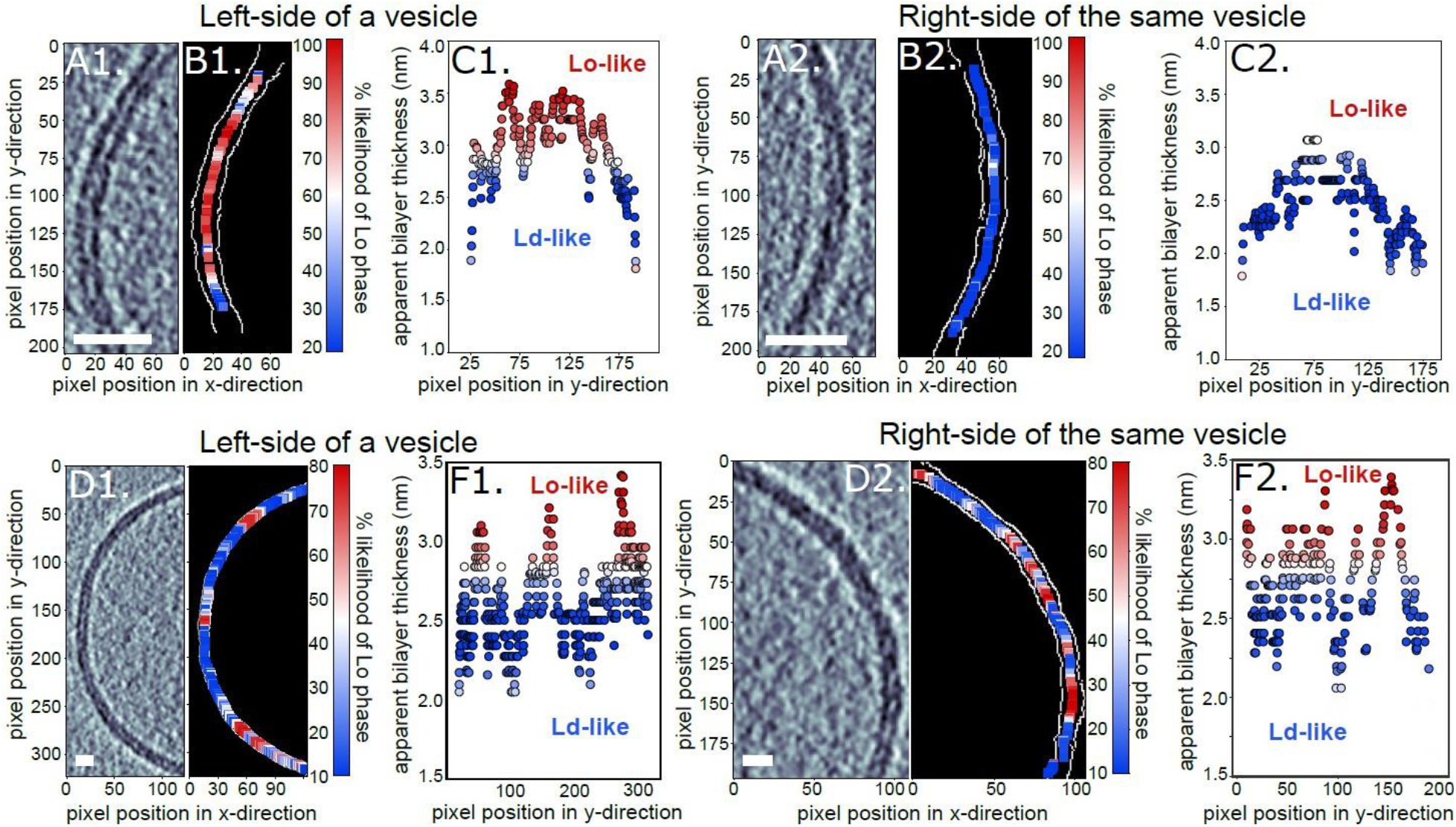
Apparent membrane thicknesses identify regions consistent with *L*_o_ vs *L*_d_ phases. Panels A, B, and C are from a single vesicle made from the lipids in Ratio 2. Similarly, Panels D, E, and F are from a single vesicle of Ratio 4. **A1, A2, D1 & D2**. Label-free tomogram slices showing bilayer regions resolved as two distinct monolayer leaflets. Scale bars are 10 nm. **B1, B2, E1 & E2**. Each slice after detection of the two edges of the bilayer (white pixels). **C1, C2, F1 & F2**. The location of each pixel on the inner edge of the bilayer vs. the minimum distance from that pixel to the outer edge. Colors represent the likelihood (from 0% to 100%) that each distance corresponds to the thicker, *L*_o_ phase instead of the thinner, *L*_d_ phase.

For the two control samples, distributions of thicknesses formed two distinguishable peaks corresponding to a thinner *L*_d_ membrane (Ratio 1) and a thicker *L*_o_ membrane (Ratio 3). We repeated this procedure for vesicles made from Ratio 2, which lies between Ratios 1 and 3.

This intermediate composition falls in a region of the phase diagram in which membranes exhibit coexisting *L*_d_ and *L*_o_ membrane phases. Because this composition is far from a miscibility critical point, domains of *L*_d_ and *L*_o_ always coarsen into micron-scale regions in taut GUVs (28). Therefore, the area fraction of *L*_d_ and *L*_o_ phases that has been previously measured in taut GUVs (24) should be equivalent to the area fraction in submicron domains imaged by cryo-ET.

In Fig. 3A, it is clear that cryo-ET of submicron vesicles of Ratio 2 indeed yields a distribution of bilayer thicknesses that corresponds to a mixture of thin and thick membranes. In Fig. 4, it is clear that this probe-free method can resolve submicron domains. Domains in Fig. 4 are constrained to submicron sizes for the obvious reason that the vesicle themselves are submicron, and perhaps also for the more subtle reason that excess area in nonspherical membranes allows submicron domain configurations (6). Next, we quantitatively evaluated the probability that each thickness corresponds to the *L*_d_ versus the *L*_o_ phase, using the mixture model described in the methods. This analysis led to the conclusion that vesicles made from Ratio 2 contain domains and that 43 ± 3% of the membrane area is in the *L*_d_ phase.

This area fraction of 43 ± 3% *L*_d_ phase, measured by cryo-ET in submicron vesicles, is in statistical agreement with values measured in vesicles that are hundreds to thousands of times larger. Quantitative tie-lines have been previously measured by NMR of multilamellar vesicles of the same lipid composition (35/35/30 DiPhyPC/DPPC/cholesterol), and edges of liquid-liquid coexistence regions have been previously measured by fluorescence microscopy of GUVs (24). These previous experiments have firmly established that micron-scale GUVs made from lipids mixed in Ratio 2 contain 50% ± 6 mole % *L*_d_ phase, which agrees with our cryo-ET values within the experimental uncertainty of the two methods.

Next, we tuned the lipid composition (and area fraction) of vesicles to show that thickness mismatches quantitatively identify domains in membranes that do not lie exactly on tie-lines, with controls that do not lie exactly at endpoints. For example, Ratios 4, 5, and 6 lie on a line that is parallel and near the tie-line of Ratios 1, 2, and 3. The two new controls (Ratios 4 and 6) are near endpoints, but are not purely *L*_d_ or *L*_o_ phases. Applying the cryo-ET imaging and analysis above leads to the conclusion that for vesicles made from Ratio 5, 83 ± 2% of the area is

*L*_d_ phase (Fig. 3D). This value is in excellent agreement with two independent measurements for micron-scale vesicles. A value of 80 ± 6 mole *%* of *L*_d_ phase is expected for intact vesicles (24), and 76 ± 6 area % of *L*_d_ phase was determined by atomic force microscopy of GUVs ruptured on mica surfaces (20). In summary, a label-free approach of identifying domains by membrane thicknesses accurately quantifies the amount of *L*_d_ and *L*_o_ phases.

Switching our focus to identify *L*_d_ or *L*_o_ domains by the second method, namely by partitioning of a probe, we surveyed labels that are both fluorescent and electron-dense. Our goal was to find a single probe to serve two purposes: to image micron-scale GUVs by fluorescence microscopy and to image submicron domains in ~100 nm vesicles by cryo-ET. This requirement of a dual use imposed several challenging criteria. The probe: **1**: must be highly electron dense so that it is visible by cryo-ET, **2**: must be fluorescent, and **3**: must partition strongly with membrane domains. Moreover, for the probe to be non-perturbing, it must meet additional criteria. **4**: Any fluorescent or electron-dense moiety must be attached to the probe through a single binding site in order to avoid over-counting (14) or crosslinking (29). **5**: The probe must partition strongly to the membrane so that it can be used at low concentrations (30). **6**: The probe must not severely perturb the membrane’s shape (as, for example, a BAR domain protein would (31)). **7:** The probe must not aggregate or induce membranes to stick to each other.

We tested a panel of 7 probes (SI Appendix, Table S1 and Fig. S3) and found that our criteria were best met by an mCherry trimer that binds through a single site to DGS-NTA(Ni) lipids incorporated into electroformed vesicles. Other probes fell short by aggregating (A206K GFP), causing vesicles to aggregate (A206K GFP), and/or producing no discernable contrast between membrane phases (14:0 PE-DTPA(Gd), GM1 lipids with Cholera Toxin B, monomeric mCherry, and 18:1 DGS-NTA(Ni) without mCherry).

In Fig. 5, we establish proof of principle that membranes labeled by a single probe can be imaged by both fluorescence microscopy and cryo-ET. To image micron-scale GUVs with our dual-use mCherry trimer, we added the probe directly to GUV solutions. Fluorescence micrographs in Fig. 5A-B show that the probe strongly preferentially partitions to the *L*_d_ phase. The absence of bright puncta in the aqueous regions implies that there is no significant protein aggregation in solution. To image smaller vesicles with our dual-use probe, we extruded unlabeled GUVs and then added mCherry trimer to the resulting solution (Fig. 1C). In many cryo-ET tomograms, clusters of mCherry trimers appear in a single layer on membrane surfaces (Fig. 5C-F), consistent with the bright labeling of domains we observed in GUVs. Trimers in the clusters are evenly spaced ~3 nm apart (SI Appendix, Fig. S4), consistent with monovalent binding to DGS-NTA(Ni) lipids in *L*_d_ domains.

**Figure 5.**
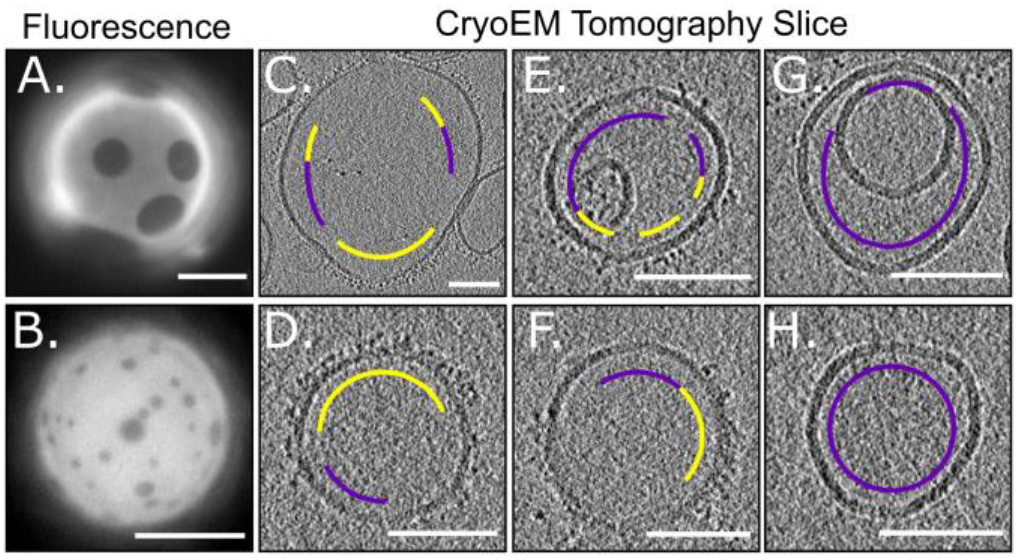
Trimeric his6-mCherry as a dual-use probe to image GUVs by fluorescence microscopy and to image submicron vesicles by cryo-ET. Lipids were mixed in Ratios 2 (top row) and 4 (bottom row). A-B. Fluorescence micrographs in which trimeric his6-mCherry labels the *L*_d_ phase of GUVs. Scale bars are 20 μm. C-H. Cryo-ET tomographic central slices of extruded vesicles. On the exterior of vesicles, regions that are densely covered by a brush of trimeric his6-mCherry (yellow arc) are clearly distinguishable from areas that are devoid of his6-mCherry (magenta arc). Scale bars are 100 nm. A larger version of this figure appears in SI Appendix, Fig. S4-S5.

The utility of any probe, including the mCherry trimer, to image domains by cryo-ET is mitigated by four observations, all of which are illustrated in Fig. 5. **1**: When vesicles touch, it is unclear if an absence of probe denotes an *L*_o_ domain or simply inaccessibility of the probe to the membrane surface. For applications in which the addition of charged lipids is acceptable, their presence can help maintain separation between membranes. **2**: When membrane regions are sparsely labeled, it is unclear whether they should be assigned to the *L*_o_ phase or to the *L*_d_ phase. **3**: Uniform mixing of the probe is challenging to achieve – some vesicles appear unlabeled (Fig. 5G-H). **4**: Two regions of the sample are unusable: the interior vesicles of multilamellar structures, which are inaccessible to the probe, and the air-water interfaces, which trap unbound probes (SI Appendix, Movie S1). An advantage of tomography is that the air-water interface can be computationally sliced away.

## DISCUSSION AND CONCLUSION

Here, we leveraged cryo-ET imaging to develop two methods of identifying liquid domains in membranes. Both methods present strengths and limitations.

The main advantage of using membrane thickness to identify liquid domains in unstained vesicles is that the technique is probe-free. It avoids all concern that labels may shift transition temperatures (30) or lead to oxidation (32). Because the analysis is statistical, it works well when distributions are built from a large number of thickness measurements. For example, the distributions of Ratios 1 and 3 in Fig. 3A reflect ~5000 points. Conversely, the method will fail if images are not representative. For example, in isolation, the micrograph in Fig. 4A2 could be misinterpreted as signifying that all membranes in Ratio 2 are nearly entirely in the *L*_d_ phase. Another feature of the technique is that it does not require vesicles to be unilamellar or spherical, so it can be applied to uncharged vesicles extruded through 100 nm pores, which are typically neither unilamellar (33) nor spherical (Fig. 2 and Fig. 5).

However, the approach is limited to membranes in which the difference in thicknesses between the *L*_d_ and *L*_o_ phases is resolvable. Luckily, many membranes fulfill this criterion (16–21). Thickness differences can be maximized through savvy choices for the types of lipids in the system and the ratios at which they are mixed. For example, mixing long, saturated lipids with short, unsaturated or methylated lipids typically results in thick *L*_d_ phases and thin *L*_o_ phases (16–20). Existing phase diagrams and tie-lines, which are reviewed in (34, 35) and discussed further in the Supplementary Information, can be leveraged to quantitatively and accurately measure the relative amounts of *L*_d_ and *L*_o_ phases.

The main advantage of using an mCherry probe to identify liquid domains is that the label is both fluorescent and electron-dense, enabling similarly prepared vesicles (made from the same batch of lipids, on the same day, using the same methods up to the final extrusion step) to be directly imaged by both fluorescence microscopy and cryo-ET. Dual-use probes of this type are necessary for some types of controls. For example, domain sizes in membranes of 13.65/25.35/39/22 DOPC/POPC/DSPC/cholesterol are reported to be different in GUVs and in ~60 nm vesicles (36); a dual-purpose probe could be used to distinguish if the discrepancy is due to different vesicle sizes or merely due to sample-to-sample variations in lipid composition. This point is powerful because lipid ratios in electroformed giant unilamellar vesicles can differ from ratios in hydrated, multilamellar vesicles (the basis of most protocols for imaging submicron domains) (37). An additional advantage of the mCherry probe is that it attached to a lipid through a single binding site, which avoids over-counting.

With all probes, it can difficult to achieve uniform labeling, especially when vesicles are near an air-water interface or are in contact with each other, as is common when vesicles are composed of only zwitterionic lipids. Likewise, for all probes, it is difficult to determine whether clusters of probes originally nucleated on the membrane or in solution. An advantage of the mCherry probe is that it has been shown to aggregate only at relatively high concentrations (>25 μM (38)). A remaining challenge is that the mCherry probe appears to strongly partition to the air-water interface of cryo-ET grids, and any protein at this location has the potential to denature. Looking to the future, productive approaches could include synthesis of quantum dots that label membranes via a single linker.

In conclusion, we have developed two new methods to identify submicron domains in membranes. One method employs thickness differences, and the other employs dual-use probes. We employ cryo-ET to generate the first direct images of <100 nm liquid domains in protein-free, model membranes under native solvent conditions. We use these images to quantitatively correlate the area fractions of *L*_d_ (and *L*o) phase in small vesicles on length scales smaller than 100 nm and in giant vesicles on length scales greater than micrometers.

Our approach complements existing methods for imaging <100 nm liquid domains in membranes, including freeze-fracture TEM (8–11), TEM of gold-labeled membranes (12, 13), AFM (15–21), and NSOM (22). All of these methods suffer from low throughput. Nevertheless, they are valuable because they circumvent limitations of spectroscopic methods (e.g. NMR, electron paramagnetic resonance, and fluorescence resonance energy transfer) and scattering methods (e.g. X-ray and neutron scattering) (36, 39–42). These indirect methods typically measure only average properties of domains and typically generate data that must be analyzed in the context of models. Direct imaging of 10-100 nm liquid domains is valuable because it can potentially allay concerns that submicron domains are artifacts (43). Direct imaging is vital for testing domain nucleation theories (44), evaluating microemulsion mechanisms (7), assessing simulations (45), and probing discrepancies between results in GUV and in ~60 nm vesicles (36).

## MATERIALS AND METHODS

### Lipids

Phosphocholine (PC) lipids (Avanti Polar Lipids, Alabaster, AL), cholesterol (chol; Sigma Aldrich, St. Louis, MO), and Texas Red dihexadecanoyl-PE (DHPE; Life Technologies, Grand Island, NY) were used as purchased without further purification. Lipid stock solutions in chloroform contained, at minimum, a ternary mixture of diphytanoyl-PC (DiPhyPC; 4 ME 16:0 PC), dipalmitoyl-PC (DPPC; 16:0 PC), and cholesterol. DiPhyPC and DPPC are zwitterionic; cholesterol is uncharged. Features of this ternary mixture is that its miscibility phase diagram has been mapped in detail (24), and the saturated carbon chains of DiPhyPC resist oxidation (5). Stock solutions for cryo-ET experiments with mCherry labels also contained 2 mol% of a nickel-chelating lipid of dioleoylglycerosuccinyliminodiacetic acid (18:1 DGS-NTA(Ni), which replaced 2 mol% DiPhyPC), whereas stocks for fluorescence microscopy controls of GUVs contained 0.8 mol% Texas Red DHPE.

### GUV electroformation

Solutions containing 2 x 10^-6^ moles (~0.76 mg) of lipids were spread evenly on slides coated with indium tin oxide. The slides were placed under vacuum for >30 min to evaporate the chloroform. A capacitor was created by sandwiching 0.3 mm Teflon spaces between two lipid-coated slides. The gap was filled with 335 mM sucrose, and the edges were sealed with vacuum grease. GUVs 10-100 μm in diameter were electroformed (37) by application of an AC voltage of 1.5 V at 10 Hz across the capacitor for 1 hr at 60°C.

### Extrusion

Two solutions were produced: a “thickness mismatch buffer” of 150 mM NaCl and 25 mM HEPES and an “mCherry buffer” of 150 mM NaCl, 25 mM HEPES, and 1 mM TCEP. GUVs were diluted 5-fold in one of the buffers and then concentrated by centrifugation at approximately 10,000 rcf for 10 min. Supernatant was removed. GUVs were re-diluted in 100 μl of buffer and stored at 60°C for <15 minutes before extrusion in order to ensure that vesicles were well above their mixing temperatures, which ranged from 25°C to 48°C. A mini-extruder (Avanti Polar Lipids) with a heat block and two 1 ml gas-tight syringes were maintained in an oven at 75°C before use. Most GUVs were extruded 29 times at 75°C through 50 nm pores in polycarbonate membranes The only exception was that GUVs of mixtures 4, 5, and 6 for thickness mismatch experiments were extruded through membranes with 100 nm pores. The resulting small vesicles were stored at room temperature for < 1 hr before vitrification.

Despite initial electroformation, extruded vesicles were often not unilamellar. This is surprising because electroformation typically yields unilamellar vesicles. We found that vesicles electroformed and extruded in pure water were always multilamellar, with at least two lamellae per vesicle. When we increased the ionic strength of the buffer with NaCl, we produced more unilamellar vesicles. Interestingly this result is the opposite to that reported in (33).

### Introduction of trimeric mCherry

Extruded vesicles containing 2 mol% DGS-NTA(Ni) lipids were diluted in “mCherry buffer”, lightly vortexed with 2 mM trimeric mCherry (46) and allowed to incubate for >25 min at room temperature. The trimer presents a larger volume of electron-dense material than a single mCherry molecule. The first mCherry in the trimer is fluorescent, and the last mCherry binds monovalently through a his6-tag to the nickel atom on a DGS-NTA(Ni) lipid (Fig. 1). At pH 7.4 (the pH of our “mCherry buffer”), the free carboxyl group on DGS-NTA(Ni) is negatively charged. By cryoET, we did not observe a difference in shape or lamellarity of vesicles with or without DGS-NTA(Ni).

### Fluorescence imaging

Immediately before imaging, GUV solutions were further diluted 10-fold in one of the buffers and sandwiched between two coverslips. The edges of the coverslips were sealed with vacuum grease. Both DGS-NTA(Ni) and Texas Red DHPE preferentially partition to the *L*_d_ phase, which appears bright by fluorescence microscopy; the *L*_o_ phase appears dark. GUV images were viewed through an air objective on a Nikon Y-FL upright epifluorescence microscope (Nikon, Melville, NY), captured on a Photometrics CoolSnapFX camera (Photometrics, Tucson, AZ), and manipulated using ImageJ (http://imagej.nih.gov/ij/). To preserve the fidelity of the data, image manipulation was limited to adjusting overall brightness or implementing linear (γ=1) contrast enhancements.

### Cryo Electron Tomography

Solutions of extruded vesicles were mixed with 6 nm colloidal gold fiducial markers (Aurion, Wageningen, Netherlands) and applied to glow-discharged C-flat holey carbon grids (Electron Microscopy Science, Hatfield, PA) or QUANTIFOIL R 2/2 holey carbon grids (Quantifoil Micro Tools GmbH, Großlöbichau, Germany) and plunge-frozen into liquid ethane using a Vitrobot Mark IV (FEI, Hillsboro, OR) at 100% humidity and temperatures of either 25°C (for height mismatch) or 4°C (for trimeric mCherry). The thinness of the water film on the grid can perturb larger vesicles by flattening them and by introducing interactions with the air-water interface. Given that membrane domains were observed in vesicles over the entire range of sizes, with expected area fractions, this perturbation is minor at the 0° plane where vesicles were evaluated.

Data were collected on a TF20 TEM (FEI, Hillsboro, OR) or a Glacios cryo-TEM (ThermoFisher Scientific, Waltham, MA) operated at 200 kV with a K2 Summit direct electron detector (Gatan, Pleasanton, CA) collecting 200 ms frames in counting mode. Frames were aligned using UCSF MotionCor2 (47) through the Appion web interface (48). Tilt-series were collected on the TF20 TEM with a bidirectional tilt-scheme of −48° to 48° in 3-degree steps at a nominal magnification of 14,500x (pixel size 2.54 Å/pixel) with a total dose of ~100 e^-^/Å^2^. Tilt-series on the Glacios TEM were collected in a dose-symmetric tilt-scheme (49) between −63° and 66° in 3-degree steps at 22,000x (pixel size 1.91 Å/pixel) with a total dose of 74 e^-^/Å^2^.

The IMOD software suite (50, 51) was used to align each tilt series and to generate tomograms with contrast transfer function correction by ctfphaseflip in Etomo with defocus estimates from CTFFIND4 (52). Resulting tomograms were binned by two to a pixel size of 5.04 Å or 3.82 Å and then median filtered. Measurements were confined to central slices of the resulting stack due to resolution anisotropy and missing-wedge effects from limited angular sampling. These slices were projected in the z-direction in groups of 10, for a composite stack 5.04 nm or 3.82 nm thick, and Gaussian filtered (ρ = 0.75) for display in ImageJ (53). To preserve fidelity of image stacks, the only contrast enhancement was linear (γ = 1). Projection images of vesicles were also collected (SI Appendix, Fig. S2), but it was challenging to achieve sufficient contrast between the leaflets of the bilayer and the background to perform thickness measurements.

### Analysis of thickness mismatches

Cryo-electron tomograms were analyzed to yield the fraction of bilayer corresponding to the *L*_o_ and *L*_d_ phases. We cropped fields of tomograms to retain only the areas in which the membrane was resolvable in the central tomographic slice as two bands of lipid headgroups with high electron density (Fig. 2C). In some cases, different lipids may give rise to different apparent electron densities due to the mass and charge of their headgroups. In Fig. 2C and Fig. 4, there is no clear evidence of two distinguishable electron densities within the bilayer regions that can be resolved as two distinct leaflets. In theory, all vesicles are spherical (or cylindrical), and the central slices of tomograms always cut perpendicularly through the equator of each vesicle, leading to sharp, distinct electron densities for each leaflet of the membrane. However, if vesicles are not spherical, then a tomogram slice can cut obliquely through part of the membrane, such that the leaflets are not resolvable at that location. An extreme example would be a vesicle in the shape of a right triangle; a thick tomographic slice that cut horizontally through the triangle could potentially resolve some features in the vertical wall of the triangle and not in the hypotenuse.

Images were Canny filtered (54) to detect the inner and outer edges of the bilayer using the original Python program “BilayerHeightMeasurements.py” that the authors have made available by public license at https://github.com/caitlin-cornell23/cryoEMliposomes. The apparent bilayer thickness was defined to be the minimum distance between each pixel on the inner leaflet and all possible pixels on the outer leaflet; this distance is not necessarily the same as the absolute bilayer thickness. It is not necessary to know the absolute thickness – the analysis identifies a difference in thicknesses between two membrane phases. A rolling average over two distance values was applied to smooth the pixel-to-pixel variation. As our study demonstrates, this method consistently resolves differences in membrane thickness on the order of ~1 nm.

Observed minimum distances, *d*, in a given sample were assigned to *L*_o_ or *L*_d_ phases using Bayesian inference. First, 20-30 images per control sample at or near the ends of tie-lines (i.e. Ratios 1, 3, 4, and 6) were used to construct Gaussian kernel density estimates (55) of the probability of observing a particular value of the distance in each membrane phase, *p*(*d*|*L*_o_) and *p*(*d*|*L*_d_), respectively. Then, for intermediate ratios of lipids (Ratios 2 and 5), the likelihood of observing a particular distance is *p*(*d*) = *p*(*d, L*_o_) + *p*(*d, L*_d_). Incorporating the fact that the phases are mutually exclusive and defining *p*(*L*_o_) as a mixing coefficient parameter yields *p*(*d*) = *p*(*L*_o_) *p*(*d*|*L*_o_) + [1 – *p*(*L*_o_)] *p*(*d*|*L_d_*). The fraction of the membrane that corresponds to the *L*_o_ phase, *p*(*L*_o_), was constrained from 0 to 1 with the otherwise uninformative prior of *p*(*L*_o_) ~ *Uniform*(0,1). This prior, in combination with the likelihood above, gives the mixing coefficient’s posterior distribution, up to a normalization constant that we determined by numerical integration. This Bayesian inference procedure is in the original Python 3.6 program “MixtureModel.py” that the authors have made available through public license at https://github.com/caitlin-cornell23/cryoEMliposomes. The mixture coefficient’s posterior distribution is described throughout the text using the mean and two standard deviations.

In Fig. 4, domains were mapped onto individual tomograms of Ratio 2 and Ratio 5 vesicles by calculating the probability that each observed distance corresponds to the *L*_o_ phase. More specifically, we used Bayes’ theorem to calculate

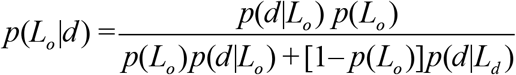

where probabilities on the right-hand side come from the calculation described above and *p*(*L*_o_) is approximated by its posterior mean. This procedure is implemented in the original Python 3.6 program “ColorMapMask_V2.py” that the authors have made available through public license at https://github.com/caitlin-cornell23/cryoEMliposomes.

## AUTHOR CONTRIBUTIONS

C.E.C., A.M., K.K.L., and S.L.K. designed the research. C.E.C. and A.M. performed experiments, collected data, and analyzed data. N.T. and C.E.C. wrote code to analyze images. C. E.C. and S.L.K. wrote the article and designed the figures.

## ACKNOWLEDGEMENTS

We thank Eva Schmid and Dan Fletcher for their kind gift of trimeric mCherry. We thank Jeanne Stachowiak for her kind gift of monomeric mCherry and A206K GFP, as well as for expert advice on the development of EM-compatible probes. We thank Avanti Polar Lipids for the gift of Aurora-DSG nanoparticles. We thank Ilya Levental and Fred Heberle for outstanding collegiality: they saw us present the results in this manuscript at the 2019 Biophysical Society Meeting, informed us of their complementary project, and simultaneously submitted with us. We hope our feedback influenced their manuscript as positively as their feedback influenced ours. Seed funding for this project was provided by UW Royalty Research Grant A122781 to S.L.K. Lipid research in the Keller lab is supported by National Science Foundation grants MCB-1402059 and MCB-1925731. K.K.L. was supported by NIH grant R01-GM099989. C.E.C. and A.M. were funded by the National Institutes of General Medical Sciences of the National Institutes of Health under award T32GM008268 (to C.E.C.) and T32-GM007750 (to A.M.).

