## Supplemental Information Appendix for "Direct Imaging of Liquid Domains in Membranes by Cryo Electron Tomography"

### Supplementary Information Appendix to "Direct Imaging of Domains in Membranes by Cryo Electron Tomography"

**CONTENTS:**

**Text S1.** Details of using tie-lines to quantify relative amounts of $L_d$ and $L_o$ phases.

**Table S1.** Probes unsuitable for distinguishing Lo and Ld phases by cryo ET

**Figure S1.** Cryo-electron tomograms of samples 2, 3, and 4

**Figure S2.** CryoEM projection images of ratios 1-3.

**Figure S3.** CryoEM tomograms and projection images of probes in Table S1.

**Figure S4.** Line scan of image from Figure 5D to determine approx. distance bound mCherry

**Figure S5.** Larger version of Fig. 5 from the main text.

**Movie S1.** Movie of reconstructed cryo-electron tomogram of the vesicle in Fig. 5E.

**References for the supplement**

**Text S1.** Details of using tie-lines to quantify relative amounts of $L_d$ and $L_o$ phases.

When quantifying relative amounts of $L_d$ and $L_o$ phases, microscopists tend to report the area fraction of each phase, whereas spectroscopists tend to report mole fractions. For precise conversions of area fractions into mole fractions, cholesterol's "area condensation" of the surrounding lipids must be considered (1). Unfortunately, area condensation values have been measured for only a small number of binary mixtures of cholesterol and PC-lipids (2–8) , and the area per lipid has been measured in an even smaller number of ternary mixtures (9).

**TABLE S1:**
**Probes unsuitable for distinguishing Lo and Ld phases by cryo ET**

**Probe:** 18:1 DSG-NTA(Ni) lipid
**Composition of vesicle:** 30/30/40 mol% diphytanoyl-PC/dipalmitoyl-PC/cholesterol
**Method of vesicle formation:** Hydration and extrusion
**How probe was incorporated:** 1 or 5 mole % in stock lipid mixture
**Problem(s):** There is no significant contrast between the Lo and Ld phases.
**Image location:** Fig. S1, Panel A

**Probe:** his-tagged A206K GFP bound to DSG-NTA(Ni) lipid
**Composition of vesicle:** 1/19/40/40 DSG-NTA(Ni)/DiPhyPC/DPPC/cholesterol
**Method of vesicle formation:** Electroformation and extrusion through 100-nm pores
**How probe was incorporated:** A206K GFP was added to vesicles after extrusion
**Problem(s):** Vesicles stick to each other, not all vesicles are labeled, and contrast is poor.
**Image location:** Fig. S1, Panel B

**Probe:** his-tagged monomeric mCherry
**Composition of vesicle:** 3/17/40/40 DSG-NTA(Ni)/DiPhyPC/DPPC/cholesterol
**Method of vesicle formation:** Electroformation and extrusion through 100-nm pores
**How probe was incorporated:** mCherry was added to vesicles after extrusion
**Problem(s):** Vesicles are successfully labeled, but contrast is poor.
**Image location:** Fig. S1, Panel C

**Probe:** Aurora™Gold-DSG Nanoparticles
**Composition of vesicle:** 5/25/30/40 Aurora™Gold-DSG/DiPhyPC/DPPC/cholesterol
**Method of vesicle formation:** Electroformation and extrusion or hydration and extrusion
**How probe was incorporated:** In vesicle formation
**Problem(s):** The nanoprobes do not remain associated with the membrane after extrusion.
**Image location:** Fig. S1, Panel D

**Probe:** Nanogold® labeled his-tag mCherry
**Composition of vesicle:** 1/19/40/40 DSG-NTA(Ni)/DiPhyPC/DPPC/cholesterol
**Method of vesicle formation:** Electroformation and extrusion
**How probe was incorporated:** Labeled mCherry with Nanogold, added labeled protein to extruded vesicles
**Problem(s):** Gold labeling was unsuccessful.

**Probe:** Gadolinium salt of DSPE (14:0 PE-DTPA(Gd))
**Composition of vesicle:** 1/29/30/40 14:0 PE-DTPA(Gd)/DiPhyPC/DPPC/cholesterol
**Method of vesicle formation:** Electroformation and extrusion or hydration and extrusion
**How probe was incorporated:** During vesicle formation
**Problem(s):** The salt dissociates in solution
**Image location:** Fig. S1, Panel E

**Probe:** GM1 and Cholera Toxin B-FITC
**Composition of vesicle:** 2/28/30/40 GM1/DiPhyPC/DPPC/cholesterol
**Method of vesicle formation:** Electroformation and extrusion
**How probe was incorporated:** Cholera Toxin B was added to extruded vesicles
**Problem(s):** The molar mass of 12 kDa is too small, and contrast was poor.
**Image location:** Fig. S1, Panel F

**Figure S1. Cryo-electron tomograms of samples 1, 2, 3, and 4.**
Slices at 0˚ through cryoET tomograms of fields of vesicles. **A.** Vesicles of Ratio 1. **B.** Vesicles of Ratio 2. **C.** Vesicles of Ratio 3. **D.** Vesicles of Ratio 4. The angular range of the tilt series in panels A and B is ~122˚. Decreased bilayer resolution at positions corresponding to 12 o'clock and 6 o'clock on each vesicle are due to missing-wedge effects (10) from limited angular sampling. Decreasing these angular ranges results in larger areas of low resolution. For example, the tomogram in panel C was collected under conditions of very limited angular range (-24˚ to 40˚) and shows large areas of low resolution (asterisks). For comparison, arrows point to several areas in which the two leaflets of bilayers are well resolved. The dark spot at the bottom of panel C is a gold bead used to align the tomogram. Figure 2C of the main text is a cropped version of panel C in this figure. Scale bars are 50 nm.

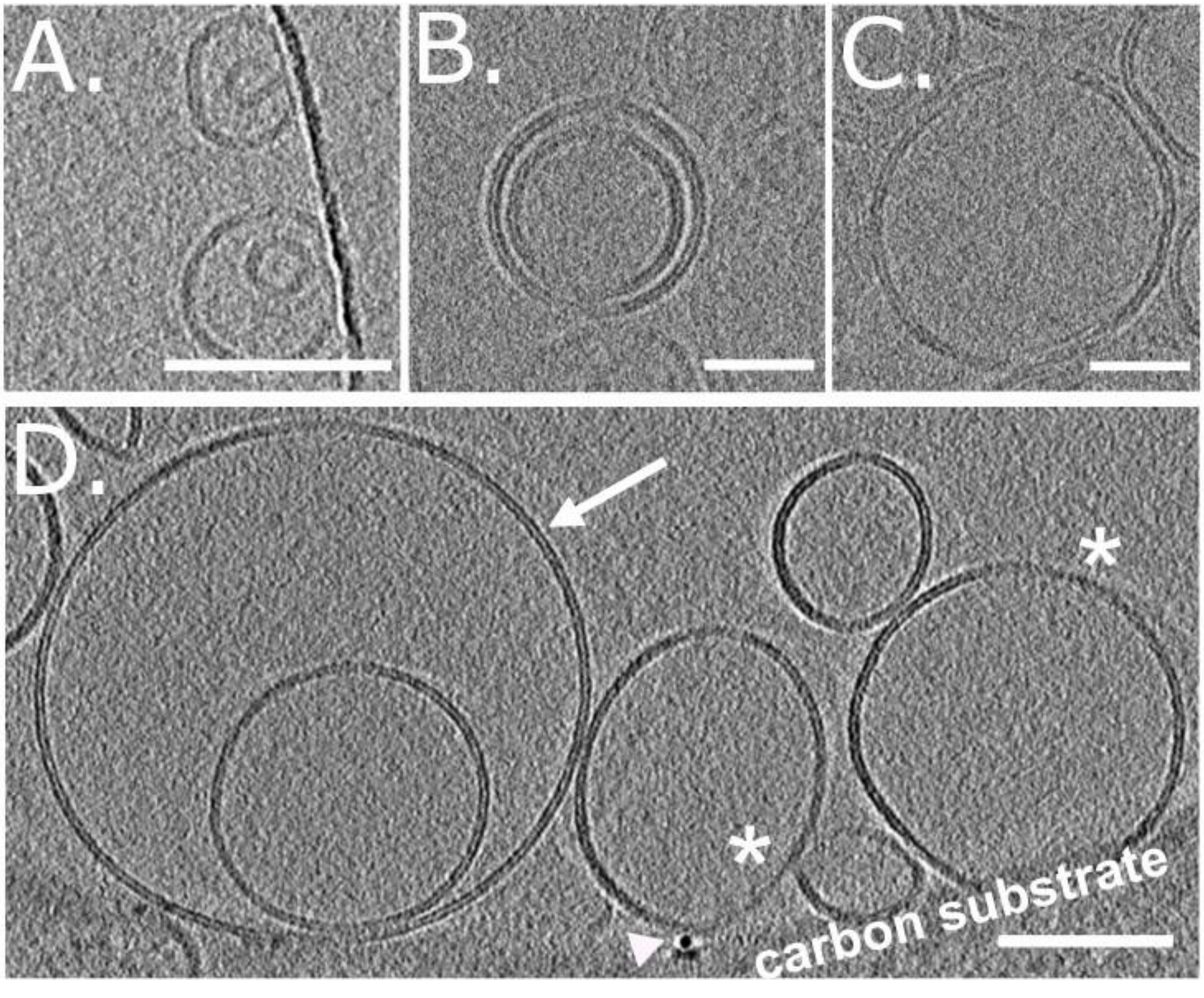

**Figure S2. CryoEM projection images of Ratios 1-3 from Table 1.**
The two leaflets of bilayer membranes are resolvable by cryoEM projection images. Dark spots in panels A and C are gold beads. Dark regions at the upper-left corner of panel A and the lower-right corner of panel B are the carbon substrate. Scale bars are 100 nm.

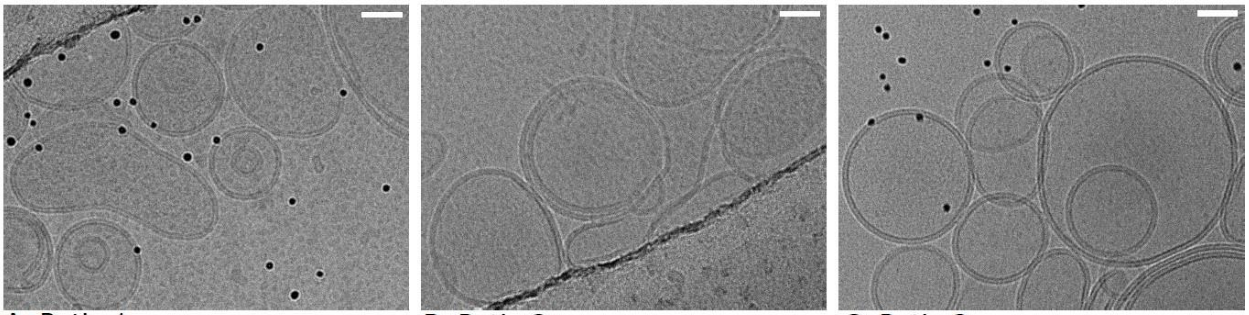

A. Ratio 1 B. Ratio 2 C. Ratio 3

**Figure S3. CryoEM tomograms and projection images of probes in Table S1.**
Images in A, B, and D-F are typical representative examples and the image in C is a "best case" example. Dark spots in panel D are gold beads to align tomograms. Dark regions in the upper-left corners of panels D-F are the carbon substrate. Scale bars are 100 nm.

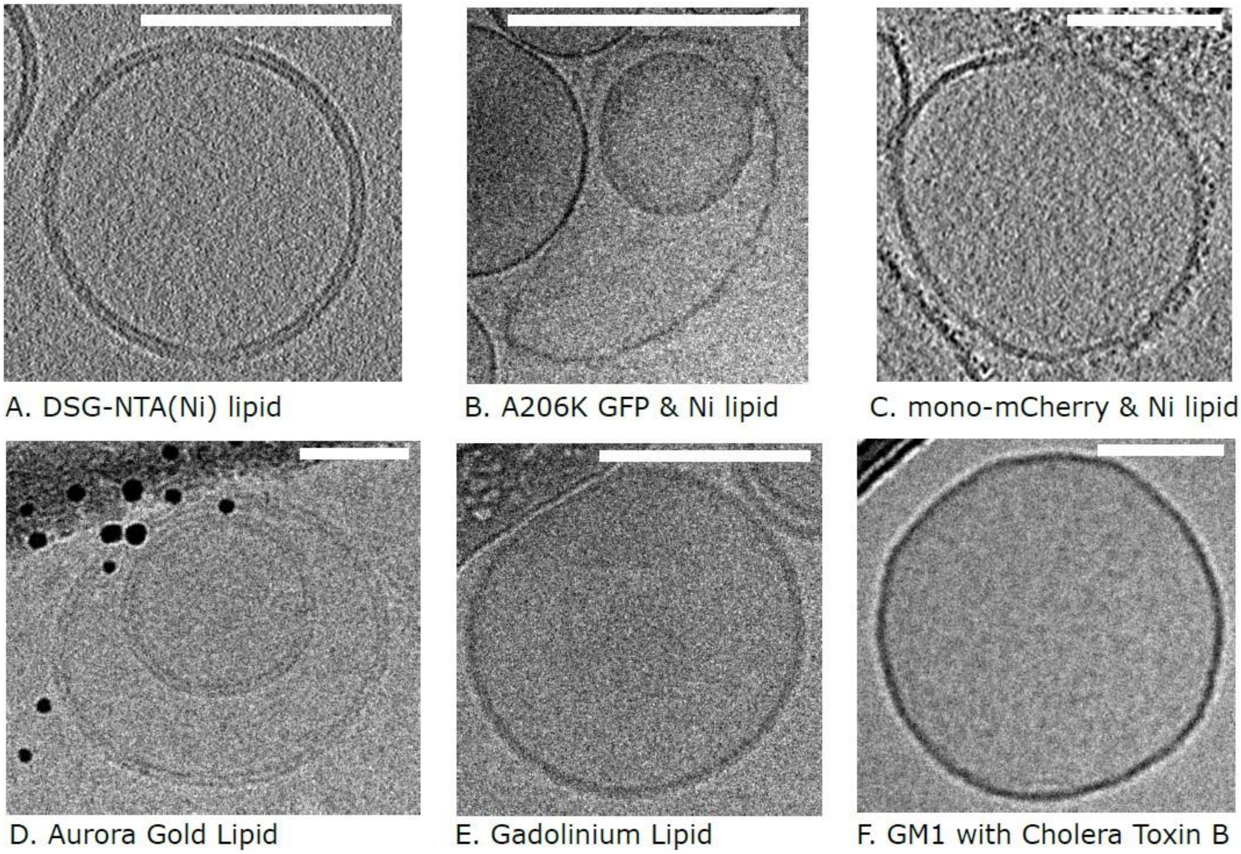

A. DSG-NTA(Ni) lipid B. A206K GFP & Ni lipid C. mono-mCherry & Ni lipid
D. Aurora Gold Lipid E. Gadolinium Lipid F. GM1 with Cholera Toxin B

**Figure S4. Intensity line scan of trimeric mCherry from Figure 5D.**
The arrow points to a region encompassing two proteins (highlighted by red lines). A line scan with a width of 5 pixels was conducted over this region, yielding the intensity profile on the right. The distance between the proteins is ~3.5 nm, the distance between the centers of the two peaks. Scale bar is 100 nm.

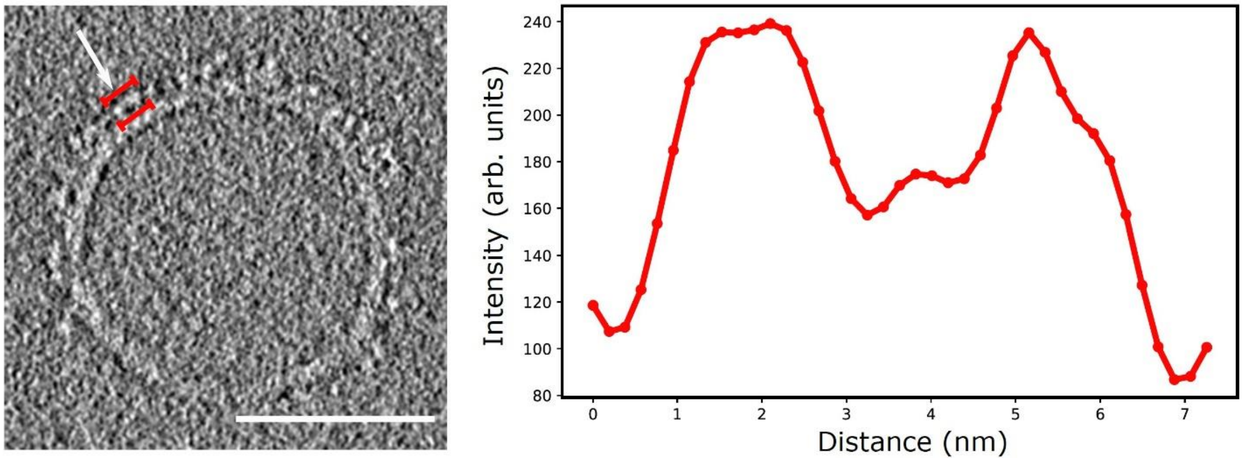


**Figure S5. Larger version of Fig. 5 from the main text.**

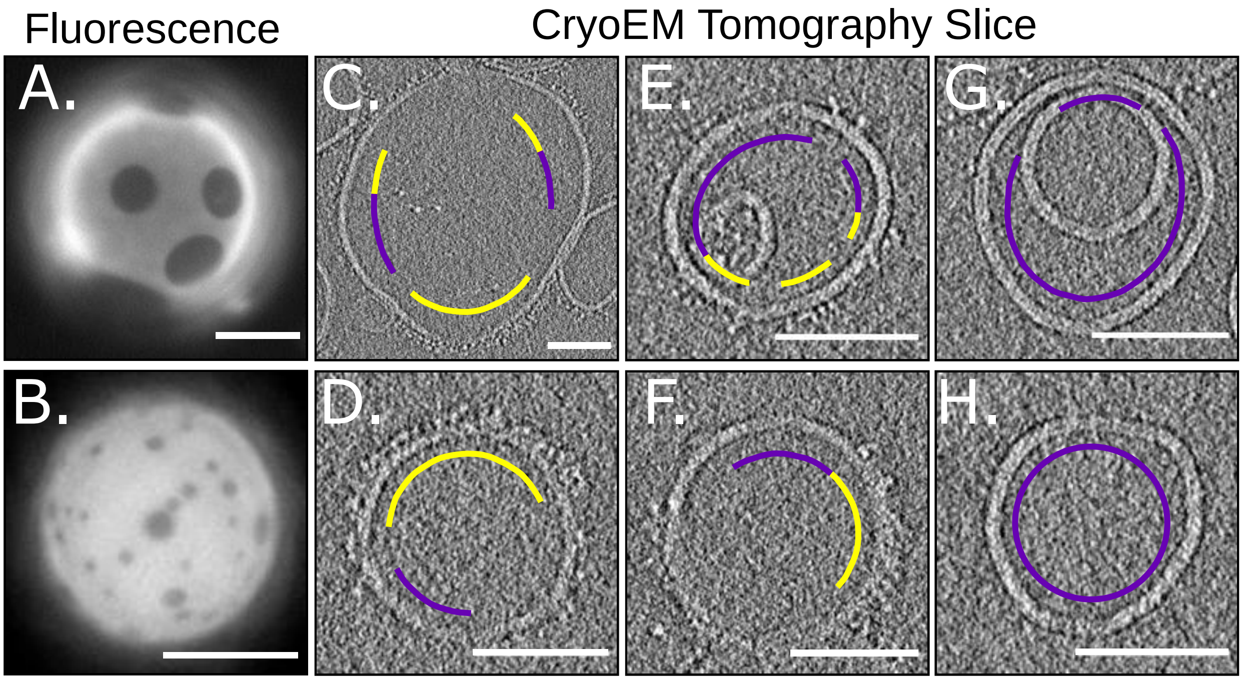


**Movie S1. Movie of reconstructed cryo-electron tomogram of the vesicle in Fig. 5E.**
The vesicle is composed of 2% DGS-Ni(NTA), 33% DiPhyPC, 35% DPPC, and 30% cholesterol. The vesicle surface has some regions that are labeled by His-tagged trimeric mCherry and some regions to which mCherry is not bound. The movie shows sequential slices in the z-axis of the reconstruction. Each slice is 3.82 nm thick. The scale bar is 25 nm.
